# Structure of the bacterial flagellar rotor MS-ring: a minimum inventory/maximum diversity system

**DOI:** 10.1101/718072

**Authors:** Steven Johnson, Yu Hang Fong, Justin Deme, Emily Furlong, Lucas Kuhlen, Susan M. Lea

## Abstract

The bacterial flagellum is a complex, self-assembling, nanomachine that confers motility on the cell. Despite great variation across species, all flagella are ultimately constructed from a helical propellor attached to a motor embedded in the inner membrane. The motor consists of a series of stator units surrounding a central rotor made up of two ring complexes, the MS-ring and the C-ring. Despite many studies, high resolution structural information is still completely lacking for the MS-ring of the rotor, and proposed mismatches in stoichiometry between the two rings have long provided a source of confusion for the field. We here present structures of the *Salmonella* MS-ring, revealing an unprecedented level of inter- and intra-chain symmetry variation that provides a structural explanation for the ability of the MS-ring to function as a complex and elegant interface between the two main functions of the flagellum, protein secretion and rotation.

---

The flagellum is the organelle responsible for the swimming motility of a huge variety of bacterial species, many of which are of clinical relevance, and the driving force behind this swimming ability has fascinated researchers since it was first observed in the 17^th^ century^1^. Flagella are highly complex, being formed from more than 25 different proteins assembled into a series of circularly symmetric and helical assemblies^2-4^. Electron cryotomographic (cryo-ET) studies have demonstrated that flagellar structures are hugely variable across species, depending on whether the flagella are to be located freely in the extracellular environment, encased in an outer membrane sheath, or entirely within the periplasm^5^. At the core of every flagellum, however, is a highly conserved inner-membrane motor that is attached to a drive-shaft, which ultimately culminates in the flagellum (Fig. 1a). The motor itself consists of a rotor complex surrounded by stator proteins that are proposed to generate torque. Rotation is rapid (up to 1,700 Hz in some species^6^), utilises ion flow through the membrane (usually a proton-motive or sodium-motive force^7^), and can respond dynamically to chemotactic signals^8^. The stators transmit the torque to a cytoplasmic complex known as the C-ring that consists of three proteins (FliG, FliM, FliN^9,10^). The N-terminal domain of FliG interacts with the extreme C-terminus of FliF^11,12^, which in turn forms the MS-ring, a large predominantly periplasmic structure that is tethered to the inner membrane via N- and C-terminal transmembrane helices^13,14^. In addition to interfacing to the C-ring to form the rotor, the MS-ring is one of the first flagellar structures to assemble^15^, and houses the type III secretion system (T3SS) of the flagellum that is responsible for the secretion and assembly of the helical components forming the drive-shaft and propellor^16^. The MS-ring therefore sits at the heart of the flagellum, both structurally and functionally. Despite this, little is known about its structure. *Salmonella enterica* serovar Typhimurium (*S*. Typhimurium) FliF consists of 560 amino acids with predicted transmembrane helices close to the N- and C-termini (Fig. 1b). Sequence analysis of residues 50-460, which lie in the periplasm, predicted that FliF consists of a series of ring building motifs (RBMs) that have previously been observed in periplasmic ring forming proteins of related secretion systems^17^. Most notably, the prediction for RBM3 was that it is formed from two disparate stretches of sequence, with a long insertion between two of the predicted β-strands. Early estimates of FliF stoichiometry from purified *S*. Typhimurium flagella suggested approximately 27 copies per flagellum^18^, and low resolution electron cryomicroscopy (cryo-EM) studies produced reconstructions with 24-,25- and 26-fold rotational symmetries applied^19,20^. Similar analyses of the *S*. Typhimurium C-ring also showed variable stoichiometry, but centred on a 34-fold symmetry^20-22^. Models of flagellar rotation have so far needed to account for these proposed mismatches in symmetry, with disagreements over the exact location of the mismatch and the functional implications (summarised in^23^. We here present near-atomic resolution structures of the MS-ring from *S*. Typhimurium that resolve these issues, revealing a conservation of stoichiometry between MS- and C-rings and unusual internal symmetry mismatches that account for the multiple functions of the rotor.

**Figure 1:**
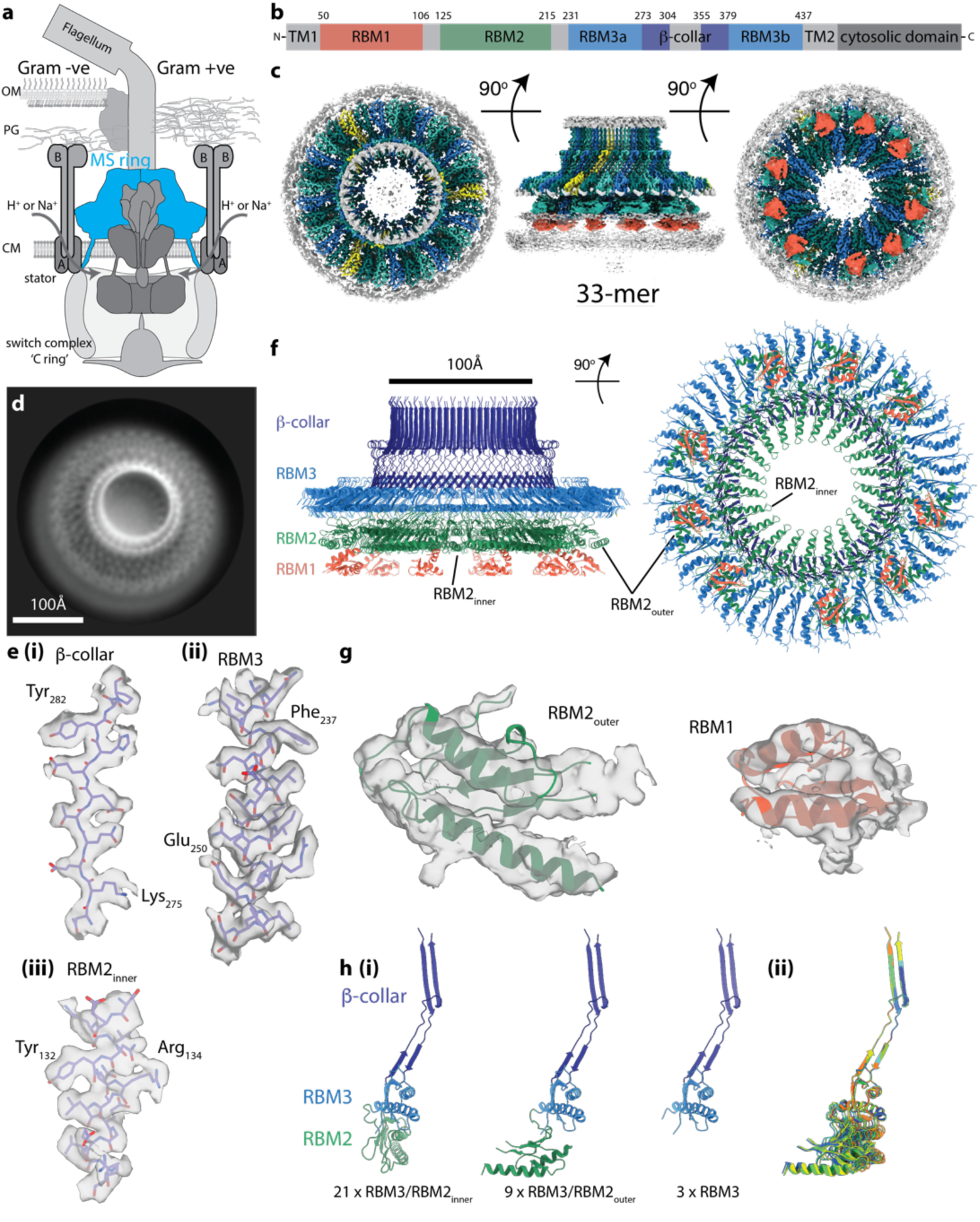
Overall structure of the flagellar MS ring. **a**, Schematic showing the location of the MS ring (blue) within the bacterial flagellum. **b**, Cartoon representation of the domain structure of FliF. **c**, Composite 3D cryo-EM reconstruction with different symmetries applied within masks (see methods). Regions occupied by RBM2 and RBM3 for each chain are similarly coloured in an alternating scheme, with the exception of chains for which only RBM3 can be seen (yellow). Regions assigned to 9 RBM1 domains are indicated in red as connectivity cannot be definitively assigned. **d**, Representative 2D class of cross-linked FliF complexes on graphene oxide surface. Scale bar, 100Å. **e**, Representative density for regions where *de-novo* building of protein domains was possible in (i, ii) the C33 averaged RBM3-region map and (iii) the C21 averaged RBM2_inner_-region map. **f**, Final model for the 33mer FliF, coloured as in (b). **g**, Representative density for docking of the RBM2_outer_ and RBM1 domains. **h**, (i) Summary of the three main conformations observed for RBM3/β-collar/RBM2 domains within the complex, coloured as in (e); (ii) overlay of the 11 copies of FliF that make up one third of the complex reveal the small changes in relative orientations of the RBM2 and RBM3 domains between different copies required to build the full object.

## FliF forms rings of mixed internal symmetry

In order to better understand how a single protein could perform multiple different roles, we over-expressed and purified the *S*. Typhimurium MS ring (FliF), using modifications of previously published protocols^13^, and collected single-particle cryoEM data from Triton X-100, DDM and amphipol A8-35 solubilised protein (Extended Data Fig. 1). Analysis of near top-down views after 2D classification (Fig. 1d) revealed a periodicity at the extremity of the largest ring consistent with > 30 subunits, rather than the expected 25-27. *Ab-initio* reconstructions and 3D-classifications were performed using a variety of imposed symmetries, but initially only a C33 reconstruction produced an interpretable map. Further refinement of this model led to a 2.6 Å volume (Fig 1c, e, f; Extended Data Table 1) in which 33 copies of residues 231-438 of FliF, corresponding to the RBM3/β-collar, were built *de novo* (Fig. 1e). However, all other densities within the C33 volume could not be interpreted as protein, suggesting either high levels of disorder or different symmetries (Extended Data Fig. 2). We therefore used the C33 particle set to perform a reconstruction in C1 which, at low resolution, revealed a periodicity underneath the C33 ring consistent with C21 symmetry. Masked refinement of this region with C21 symmetry imposed led to a 2.9 Å reconstruction (Fig. 1c, e, f; Extended Data Table 1) in which residues 125-222, corresponding to RBM2, could be built. Further refinement of these particles imposing the common C3 symmetry of the two main rings revealed density consistent with a further nine copies of RBM2 decorating the outside of the 21-fold symmetric RBM2_inner_ ring (Fig. 1g). Underneath each copy of RBM2_outer_ is a smaller density with secondary structures features consistent with a homology model for RBM1 (Fig. 1g). The overall visible structure therefore contains twenty one copies of FliF contributing one RBM3 to the 33-fold ring and one RBM2_inner_ to the 21-fold ring, nine copies of FliF contributing one RBM3 to the 33-fold ring and one RBM2_outer_, and three copies of FliF only contributing one RBM3 to the 33-fold ring (Fig. 1h). In addition we observe density consistent with nine copies of RBM1 and, although we assume they pair with the nine copies of RBM2_outer_, we cannot see linking residues to confirm this connectivity. We see no clear protein density that accounts for the remaining three RBM2s, the remaining twenty four RBM1s, or any of the transmembrane helices and cytoplasmic portions, but we found no clear evidence of proteolytic fragments in gels (Extended Data Fig. 1) or by proteomic analyses (data not shown). We did however observe further diffuse densities including a ring of material close to the observed C-terminal residues, which is consistent with detergent micelle, and a column of weak density in the centre of the structure underneath the RBM2_inner_ ring (Extended Data Fig. 3).

**Figure 2:**
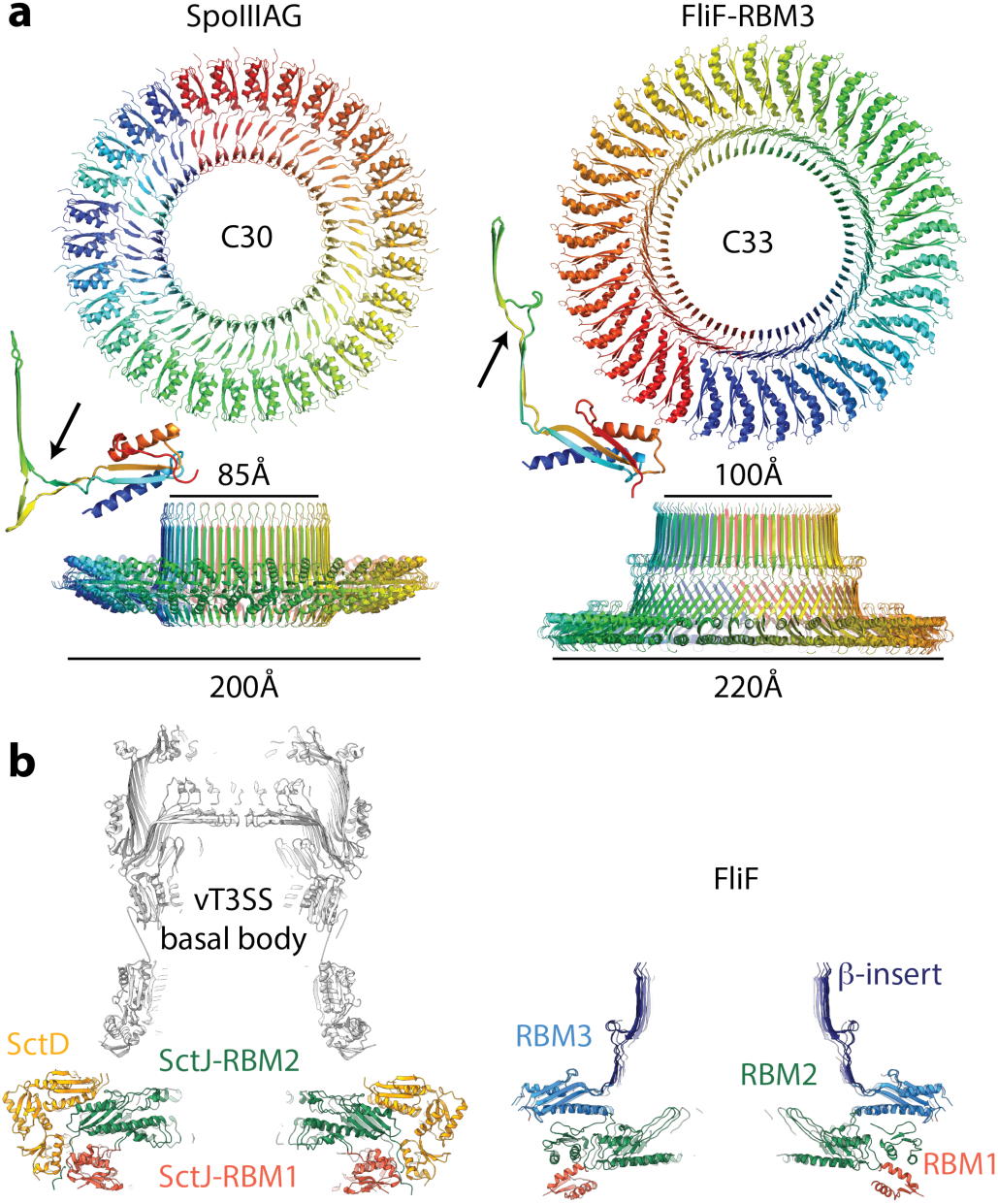
Comparison to structurally or functionally homologous assemblies. **a**, The closest structural homologue to the RBM3 portion of the FliF ring is the 30-mer fungal protein SpoIIIAG (PDB-5wc3). A view from the outer-membrane side is shown above and from the side below, with a cartoon representation of a single, extracted, monomer also shown. A small beta-insertion structure is indicated (arrow on monomer structures). **b**, The virulence T3SS basal body is constructed from two protein chains in the MS-ring equivalent region, which both form 24-mer rings consisting of multiple RBM domains. Central sections of the Salmonella SPI-1 injectisome basal body (PDB-5tcr, LH panel) and the FliF ring (RH panel) show the striking similarity in overall shape despite fundamental differences in the chains and domain types used. They also show the 21-fold RBM2_inner_ domains are very similarly arranged to the PrgK/SctJ-RBM2 24-fold ring, whilst the FliF-RBM1 and PrgK/SctJ-RBM1 domains are very differently arranged with respect to these.

**Figure 3:**
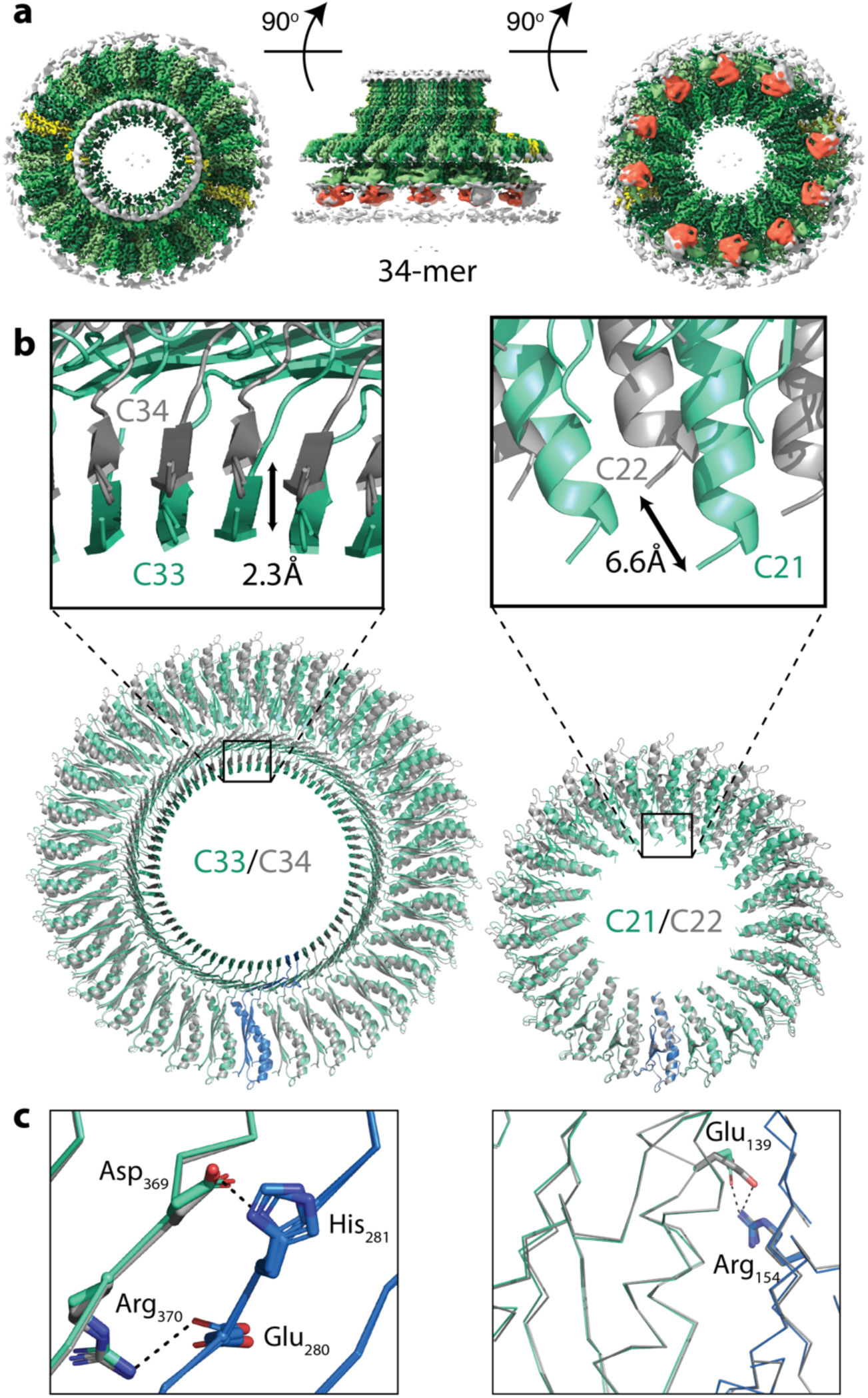
The flagellar MS ring is structurally heterogenous. **a**, Composite 3D cryo-EM reconstruction from a 34-fold stoichiometric subset of particles. C34 symmetry is applied within the RBM3 region, C22 within the RBM2_inner_ region and C2 symmetry applied elsewhere. The colour scheme mimics that of Figure 1c. **b**, Comparison of the C33/C34 and C21/C22 regions by overlaying the complete rings using a single chain reveals the subtle differences in the sizes of the respective ring-like assemblies built. **c**, Despite assembling to form rings of different symmetries, the specific interactions from which they are built are entirely conserved, including salt bridges, in both the C33 (cyan and blue)/C34 (grey) rings (left hand panel) and the C21 (cyan and blue)/C22 (grey)rings (right hand panel).

## FliF monomer structures

The enormous complexity of the 33-fold MS-ring means that there are a variety of monomer structures, with each of the 11 chains in the nominal asymmetric unit being unique in terms of relative domain orientation (Fig. 1h). Each chain, however, is made up of equivalent domains that match the predicted structural arrangement well (Fig. 1b). The density that corresponds to RBM1 fits a homology model based on domain 1 of the type III secretion system (T3SS) injectisome protein SctJ, with a βαββα topology. RBM2 and RBM3 are both canonical RBM domains with an αββαβ topology. Despite sequence identity of only 22%, they are structural homologues (rmsd of 2.3 Å over 78 Cα) (Extended Data Fig. 4). RBM2 is most closely related to domains from SctD (rmsd of 2.2 Å over 85 Cα) and SctJ (rmsd of 1.1 Å over 79 Cα), the injectisome basal body proteins that form 24-fold symmetric concentric rings tethered to the inner membrane^24^ (Extended Data Fig. 5). RBM3 on the other hand is a closer structural homologue of the RBM domain from SpoIIIAG (rmsd of 2.3 Å over 80 Cα), a sporulation protein from *Bacillus subtilis* that forms 30-fold symmetric rings in the periplasm^25^ (Extended Data Fig. 6). Both the SpoIIIAG RBM domain and RBM3 of FliF contain a long β-strand rich insertion between the first two β-strands of the RBM fold. The β-insertion (residues 273-379) essentially forms a pair of 2-stranded, anti-parallel β-sheets, one angled ∼60° from the horizontal followed by an unusual vertical section. Residues 305-354 at the tip of the vertical strands are not observed, consistent with predictions of disorder in this region due to a high number of Pro, Ser and Thr residues. A prominent loop (residues 284-292) between the sections means that the strands cross over and their relative positioning is swapped between the angled and vertical sheets (Extended Data Fig. 7). The C-terminus of RBM3 is the last observed residue in the structure and is directed towards the detergent micelle.

**Figure 4:**
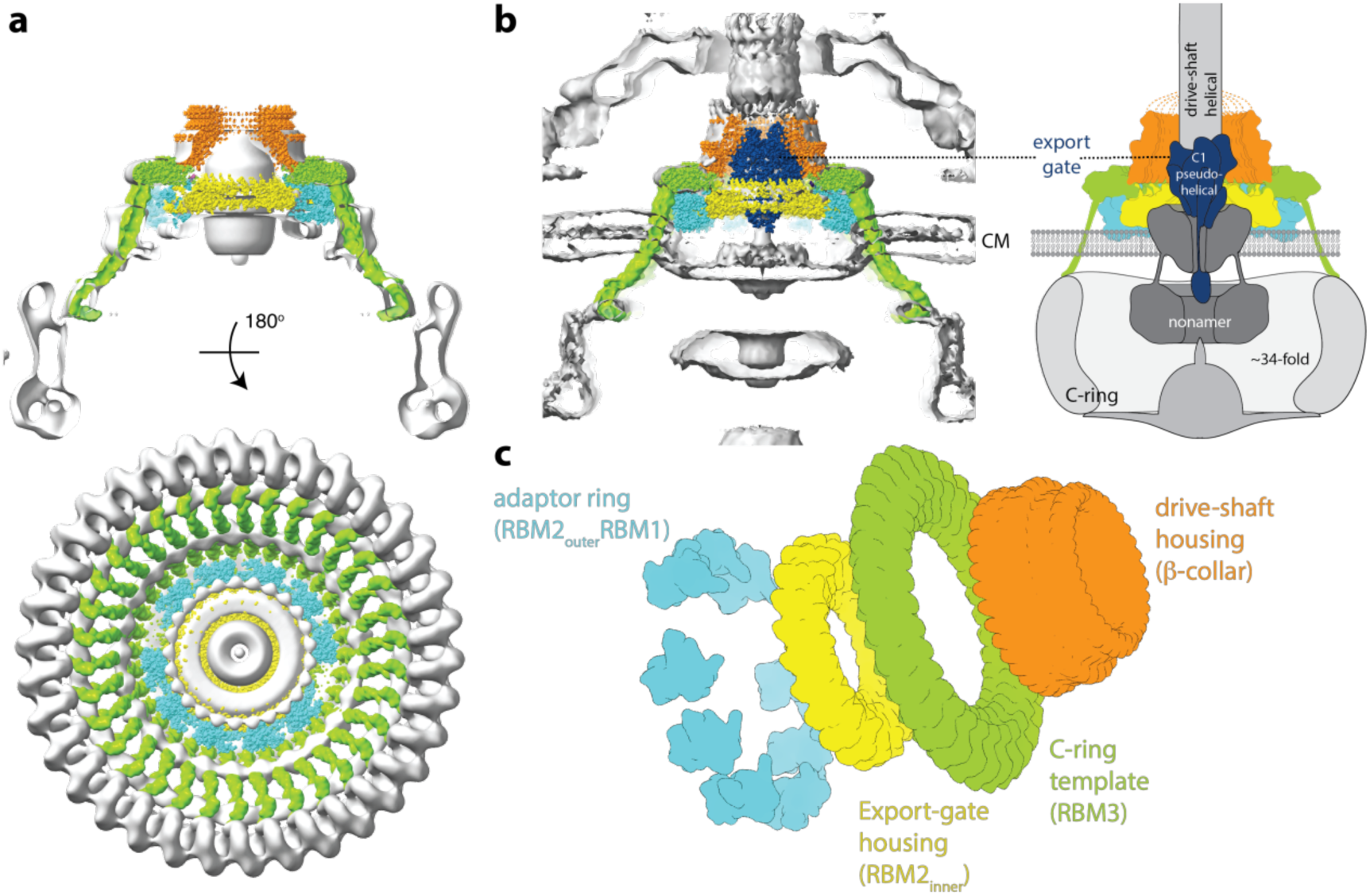
The MS-ring as a structural adapter. **a**, A model for the 34-mer MS-ring, coloured to highlight the different structural regions, is placed in the single particle reconstruction of the *S*. Typhimurium flagellar basal body (grey) (EMD-1887), showing the good match in overall shape and links to the 34-fold symmetric C-ring. The 34-mer FliF was built in the map shown in Figure 3a and extended to the C-terminus using a continuous helix of the correct length, ending in a homology model based on the crystal structure of residues 523-559 of *Helicobacter pylori* FliF (PDB: 5wuj). **b**, The FliF model (coloured as in (a)) is shown placed in a *P*. *shigelloides* tomographic volume (EMD-10057) and a model for the export gate complex (blue) (PDB-6r69) is then docked within FliF. The panel on the right is an update of the cartoon from Figure 1a, using this colour scheme. **c**, Exploded diagram of FliF coloured to emphasise the roles the different symmetries play in adapting between components within the flagellar assembly.

Overlay of all of the monomers in the structure based on the RBM3/β-collar domains highlights the structural complexity of the rotor (Fig. 1h). The RBM_inner_ and RBM2_outer_ positions are related by a 120° rotation and significant changes in the linker between RBM2 and RBM3 are required. However, even the positions of each RBM2_inner_ domain display relative rotations of up to 9° in order to accommodate the symmetry mismatch between the 33-fold and 21-fold symmetric rings.

## Assembly descriptions

The total surface area buried in the FliF ring is enormous (217000 Å^2^), totalling 36 % of the available monomer surface. All of the interaction surfaces observed in the assembly are highly conserved, with areas of greatest variation occurring in surface loops and disordered regions (Extended Data Fig. 8). Analysis of the electrostatic surface potential of the monomers reveals that the interaction surfaces are mostly hydrophobic, but patches of complementary charge are observed (Extended Data Fig. 9).

The complex can be broken down into four main structural assemblies: the 33-fold symmetric β-collar, the 33-fold symmetric RBM3 ring, the 21-fold symmetric RBM2_inner_ ring and the decorating RBM2_outer_/RBM1 domains. The β-collar accounts for 77000 Å^2^ of the buried area and consists of 66 vertical β-strands (shear number of 0) linked to 66 β-strands angled ∼60° from the horizontal. In addition to the standard β-sheet hydrogen bond network, this sub-structure is stabilised by numerous sidechain mediated hydrogen bonds and two potential salt bridges (His281-Asp369 and Glu280-Arg370). In addition, we observed a density connecting Arg373 and Lys275 from neighbouring subunits, consistent with a glutaraldehyde cross-link formed during the final purification stage before imaging (Extended Data Fig. 10).

Both of the RBM3 and RBM2_inner_ rings are constructed from the type of interface observed in other secretion system ring-forming motif structures, with the helices of one domain packing against the β-sheet of the neighbouring domain. However, when comparing the two interfaces, there is an ∼6.5° rotation of one domain relative to the other in order to accommodate the different stoichiometries of the rings (Extended Data Fig. 11). The RBM3 and RBM2_inner_ rings are further stabilised by Glu242-Arg248 and Arg154-Glu139 inter-subunit salt bridges respectively (Fig. 3c). With the exception of the linkers between them, there is virtually no contact between the RBM3 and RBM2_inner_ rings (Extended Data Fig. 12). This is likely a reflection of the fact that the symmetry mismatch between the rings both prevents a consistent interaction surface and leads to a significantly smaller diameter for the RBM2_inner_ ring (70 Å) compared with the RBM3 ring (140 Å) or the β-collar (100 Å). Contacts between the two mis-matched rings are instead via the RBM2_outer_ subunits. The C-terminal loop of one RBM2_outer_ subunit tucks between two RBM3 subunits in the ring above, while the second α-helix of the αββαβ motif bridges two RBM2_inner_ subunits rings (Extended Data Fig. 12). Additional contacts are observed between the C-terminus of an RBM1 domain and two of the RBM2_inner_ subunits. However, the lower resolution of these portions of the structure suggests these are not tight contacts.

The mechanism by which such a complex arrangement of subunits and mixed symmetries could be built from a single protein chain is intriguing. It appears that the interaction surfaces of the RBM3 and RBM2 domains are primed to build rings of significantly different stoichiometry, and hence there is a need to build in flexibility and the extreme symmetry breaking innovation of the RBM2_outer_ conformations. The majority of the structure is built from units containing two copies of the RBM2_inner_ conformation and one copy of the RBM2_outer_ conformation, presumably driven by limitations of the conformations the linkers can take preventing more than two consecutive RBM2_inner_ conformations. These blocks could either be visualised as an RBM2_inner_/RBM2_outer_/RBM2_inner_ arrangement, in which the RBM2_outer_ subunit bridges the two RBM2_inner_, or as an RBM2_inner_/RBM2_inner_/RBM2_outer_ arrangement, in which case the RBM2_outer_ provides the bridge to the next unit. This pattern is observed for three such units (contributing 9 RBM3s to the 33-fold ring and 6 RBM2_inner_s to the 21-fold ring), at which point the pattern is broken by a subunit pair containing one copy of an RBM2_inner_ conformation and one copy for which there is no visible RBM2 density. This completes one third of the structure and this pattern of packing is then repeated twice more.

## FliF combines elements of sporulation and secretion system structures

The closest structural homologue of the 33-fold RBM3 domain assembly is the SpoIIIAG protein from the sporulation system of *B*. *subtilis*^25^, which forms a 30-fold symmetric structure utilising a very similar interaction surface to the RBM3 domains of FliF (Fig. 2a and Extended Data Fig. 13). Strikingly, SpoIIIAG also contains a β-insertion that forms a 60-strand β-collar with a shear number of 0, and both proteins also share a feature of a short triangular insertion at the point the strands change direction to the vertical (Fig. 2a). Such vertical β-strands are highly unusual, but have also been observed in the outer membrane secretin structures of T3SS and type II secretion systems (T2SS)^24,26^. Despite the strong similarities, there are significant differences between FliF and SpoIIIAG, most notably in the angle of the RBM domain to the β-collar (Fig. 2a, Extended Data Fig. 13).

The RBM2 domain is most closely related at both sequence and structural levels to the RBM2 domain of the SctJ family from the virulence T3SS injectisomes and again this homology extends to the ring structures formed (Fig. 2b). The inner-membrane proximal portion of an injectisome basal body is formed by two different proteins, SctD and SctJ. The inner SctJ ring has been shown to house the export gate structure of the secretion system in its central cavity^27,28^ and forms a 24-fold symmetric ring from RBM1 and RBM2 domains^24^. The interaction interface between neighbouring domains in the SctJ RBM2 ring is closely related to the FliF RBM2_inner_ packing interaction (Extended Data Fig. 14), although the copy number difference does lead to a small difference in the size of the cavity.

## FliF exists in multiple stoichiometries

More detailed analyses of 3D classifications of the FliF particles imposing C3 symmetry revealed a subset of particles in which the C33 features were subtly broken. Further classification of these particles in C1 only allowing local angular sampling revealed that they corresponded to a C34 symmetric MS-ring. Refinement of this volume with C34 symmetry imposed led to a 2.8 Å structure of the RBM3/β-collar region of this form of the MS-ring (Fig. 3a). Initial attempts to reconstruct the RBM2_inner_ region of these particles with C21 symmetry were unsuccessful and so the C34 reconstruction was used as a reference in a C1 reconstruction that revealed 22-fold symmetry. Masked refinement of the RBM2_inner_ region with C22 symmetry imposed led to a 3.1 Å structure of this portion, while refinement of the whole volume with the common C2 symmetry led to a 3.3 Å reconstruction (Fig. 3a and Extended Data Fig. 15). Surrounding the 22-fold symmetric RBM2_inner_ ring we observed ten densities that correspond to the RBM2_outer_/RBM1 domain pairs observed in the 33mer structure, but again no density was observed for the final two copies of RBM2 or for twenty four copies of RBM1.

The MS ring is therefore capable of assembling into rings of differing stoichiometries. Analysis of the interfaces buried in the FliF 34mer revealed that only very subtle changes are needed to build the alternate stoichiometry (Fig. 3b, c). The interfaces used in the 33-fold RBM3/β-collar ring are identical to those in the 34-fold RBM3, maintaining all of the bonding interactions, including the salt bridges (Fig. 3c). A similar pattern is observed for the RBM2_inner_ rings. Although the changes are subtle, when propagated around the number of copies in the ring, they do make a difference to the diameter of each ring, with a 2.3 Å (2 %) increase seen in the RBM3/β-collar ring and a 6.6 Å (9 %) increase observed for the RBM2_inner_ ring. The most significant difference between the two structures exists in the RBM2_outer_/RBM1 domain pairs, where an extra copy is observed. However, the mode of packing of the RBM2_outer_/RBM1 against the RBM2_inner_ ring and the basic 2:1 (RBM2_inner_:RBM2_outer_) building block is conserved. The larger rings permit five copies of the trimer building block to assemble before the pattern is broken by the minority 1:1 (RBM2_inner_:RBM3only) block. It is worth noting that the symmetries of the two main rings always leave twelve copies of the monomer which don’t contribute to RBM2_inner_ and twenty four copies of RBM1 for which we see no density, although the significance of these observations is currently unclear.

Once we had observed two different assemblies in our sample, we attempted to assess whether other symmetries were also present at lower levels. To achieve this we performed supervised 3D classifications in C1 using reference models generated to reflect RBM3 symmetries from C32 to C36 (Extended Data Fig. 16). This analysis confirmed that the majority of the particles partitioned into the C33 and C34 classes (40% and 23% respectively), with 7% and 9% ending up in the C32 and C35 classes respectively. The remaining 20% went into the C36 class, but reconstructions of these particles were very low resolution and clearly artefactual.

## The MS-ring as structural adaptor

The structural heterogeneity observed in this study may seem surprising for a core component of such a fundamental cellular structure, but agrees with earlier demonstrations of stoichiometric heterogeneity for the *S*. Typhimurium C-ring^20-22^. The C-ring is a large cytoplasmic structure that assembles on to the MS-ring via a mechanism in which the N-terminal domain of the first C-ring protein, FliG, folds around two helices at the C-terminus of FliF^11,12^. The other domains of FliG then recruit the other C-ring components, FliM and FliN^29-32^, as well as providing the interaction surface for the stator complexes that generate torque^33,34^. The MS-ring/C-ring junction is therefore critical for flagellar function. The large diameter (∼ 450 Å in *S*. Typhimurium) and strong periodicity led to robust estimates of C-ring stoichiometry in both fully assembled flagella and in reconstituted MS-ring/C-ring structures. These studies revealed clear stoichiometric heterogeneity, with subunit numbers ranging between 32 and 36 copies^20-22^. The apparent mismatch between this and the originally proposed 25/26-fold symmetry of the MS-ring were confusing, especially in light of the co-folding of MS-ring and C-ring structures, and had led to models whereby symmetry mismatch at some point between the two rings was important for function.

Our structures of FliF demonstrate that the stoichiometry of the MS-ring and the C-ring are likely matched, suggesting that the entire C-ring stoichiometry is nucleated by the stoichiometry of the MS-ring. Fitting of FliF into the only available structure of a purified flagellum with an intact C-ring^20^ demonstrates the perfect fit of the dimensions of the object within the MS-ring portion of the volume, despite this region of the volume being averaged with 25-fold symmetry (Fig. 4a). Although we do not observe the C-terminal residues of FliF in our structure, the positioning of the RBM3 domains on the outside of the ring mean they are correctly placed to reach down to the FliG ring underneath the membrane. This observation was confirmed by placing the FliF structure into a subtomogram average of *in situ* flagella from *Plesiomonas shigelloides* (Fig. 4b)^35^. Interestingly this placement also provides further insights into other roles the symmetric complexity of the MS-ring may play in acting as a single chain structural adaptor molecule at the centre of the system (Fig. 4c). The RBM3 domains of the structure, and hence the cytoplasmic C-termini, have the 33/34-fold symmetry required to assemble the C-ring. The RBM2_inner_ domains, on the other hand, form the 21/22-fold symmetric ring that is seen to house the export gate in the homologous injectisome structures^24,27,36^. The highly conserved dimensions of the export gate^37^ compared to the large diversity in C-ring size between bacterial species drives the requirement for symmetry mismatch between the different domains of FliF. The subtle differences in size between the central pore of the RBM2_inner_ 21/22mers and the equivalent 24mer SctJ injectisome ring suggests there is some flexibility in the details of how the export gate is accommodated, perhaps related to the differences within the inner membrane region located below this ring seen when comparing cryo-ET of flagella and injectisomes^38^. In both systems a nonameric protein complex, termed the FlhA ring in flagella, forms a cytoplasmic ring directly below the basal body, with its transmembrane domains presumed to occupy the membrane underneath the RBM2 ring^36,39,40^. It is noteworthy that a mutation in the β-sheet of the RBM2 domain of FliF (deletion of residues 174 and 175) can be suppressed by secondary mutations in the TM domains of FlhA^41^, suggesting some ability for changes in the stability of one ring to be compensated by changes in the other. At the other side of the FliF assembly, mutations within the disordered loop at the top of the β-collar (Asn318) weaken interactions with the flagellum, and revertant mutations map to components of the proximal rod that forms the flagellar drive-shaft^42^.

## Conclusion

This study has provided, for the first time, a near-atomic resolution view of the MS-ring of the bacterial flagellar rotor. The structures reveal unexpected symmetries and an unprecedented level of structural heterogeneity for a homo-oligomeric assembly. The symmetry mismatches within the structure demonstrate how the MS-ring is able to bridge multiple different structural and functional units within the flagellar basal body utilising a single protein chain (Fig. 4c). The explicit linking of the MS-ring stoichiometry to that of the C-ring introduces new questions of how rotors of different sizes in different species of bacteria can be reconciled with this model, especially given the constraints that the need to house the T3SS in the centre of the structure places on the system. Will FliF provide yet more surprises or will other adaptor proteins play a role?

## Acknowledgements

We thank E. Johnson and A. Costin of the Central Oxford Structural Microscopy and Imaging Centre for assistance with data collection. H. Elmlund (Monash) is thanked for access to SIMPLE code ahead of release. We thank Morgan Beeby (Imperial College) for access prior to publication, to the *P*. *shigelloides* tomographic volume. The Central Oxford Structural Microscopy and Imaging Centre is supported by the Wellcome Trust (201536), The EPA Cephalosporin Trust, The Wolfson Foundation and a Royal Society/Wolfson Foundation Laboratory Refurbishment Grant (WL160052). Work in SML’s lab is supported by Wellcome Trust Investigator (100298) and Collaborative awards (209194) and an MRC Programme Grant (MR/M011984/1). L.K. is a Wellcome Trust PhD student (1009136).

## Materials & Methods

Chemicals were from Sigma-Aldrich unless otherwise specified. Detergents n-dodecyl-maltoside (DDM), Lauryl Maltose Neopentyl Glycol (LMNG) and amphipol A8-35 were from Anatrace.

### Protein expression

The FliF expression plasmid was designed based on the pKOT105 plasmid from Ueno *et al*^13^. Briefly, the *fliF* gene from *Salmonella enterica* serovar Typhimurium was amplified using Q5 polymerase (NEB) and inserted into the BamHI site of pET-3b (Merck) using NEBuilder HiFi Master Mix (NEB). FliF was expressed in *Escherichia coli* BL21 (DE3) pLysS. 20 ml of overnight culture grown at 37 °C was used to inoculate 2 L of LB media, grown at 37 °C until OD_600_ reached 0.5 and induced with 0.5 mM IPTG at 30 °C for 4 hours. Cells were harvested by centrifugation at 5000 x g for 10 minutes and frozen at −20 °C until use.

### Protein purification

Frozen cell pellet was resuspended in 40 ml of lysis buffer (50 mM Tris pH 8, 50 mM NaCl, 5 mM EDTA) and lysed by 3 passes through an Emulsiflex C5 homogeniser (Avestin) at 10,000 psi. After centrifugation at 20,000 x g for 20 min to remove cell debris, cell membranes were collected by ultracentrifugation at 186,000 x g for 1 hour. Collected membranes were dissolved in 40 ml of alkaline buffer (50 mM CAPS pH 11, 5 mM EDTA, 50 mM NaCl, 1 % (w/v) DDM) at 4 °C for 1 hour. Undissolved material was removed by centrifugation at 20,000 x g for 20 minutes. Solublised FliF was then pelleted by ultracentrifugation at 143,000 x g for 1 hour. Pelleted FliF was resuspended in 2 ml of resuspension buffer (25 mM HEPES pH 8, 50 mM NaCl, 0.1 % (w/v) DDM). FliF ring assemblies were then separated from FliF monomers by loading the resuspended FliF on a 15-40 % (v/v) sucrose gradient prepared using gradient buffer (10 mM HEPES pH 8, 5 mM EDTA, 0.02 % (w/v) DDM). The gradient was then centrifuged at 25,000 rpm for 15.5 hours using SW55Ti rotor. Gradient fixation (GraFix^43^) was used to improve stability of FliF ring assembly by addition of a 0-0.2 % (v/v) glutaraldehyde gradient to the sucrose gradient. Selected fractions of the sucrose gradient containing FliF ring assemblies was dialysed against dialysis buffer (25 mM Tris pH 8, 50 mM NaCl, 0.02 % (w/v) DDM) overnight to remove sucrose and concentrated to the appropriate concentration using a 300 kDa MWCO concentrator. For FliF preparations using Triton-X100, the same concentration of Triton-X100 was used in place of DDM.

Amphipol trapping of GraFix crosslinked FliF purified in DDM was performed by addition of amphipol A8-35 to 0.8 mg/ml FliF at 1:3 (w/w) ratio. Excess detergent was removed by the addition of BioBeads (BioRad) at 20-fold excess of detergent mass. Excess amphipol was removed by buffer exchanging into detergent-less dialysis buffer using a 100 kDa MWCO concentrator.

### Cryo-EM sample preparation and imaging

FliF samples were added to 300 mesh R1.2/1.3 Quantifoil Cu grids coated with graphene oxide substrate, blotted using Vitrobot Mark IV (FEI) and frozen with liquid ethane. The grids were imaged using a 300 keV Titan Krios microscope (FEI) with an energy filter and Gatan K2 detector (Gatan). Data were collected with a pixel size of 0.822 Å and an exposure of 1.5 e/ Å^2^/frame for 32 frames. For the sample in Triton X-100, 6111 movies were collected. For the sample in DDM, 9173 movies were collected. For the sample in amphipol A8-35, 11538 movies were collected.

### Cryo-EM data processing

Micrographs were initially processed in real time using the SIMPLE pipeline^44^, using SIMPLE-unblur for motion correction, SIMPLE-CTFFIND for CTF estimation and SIMPLE-picker for particle picking. Following initial 2D classification in SIMPLE to remove poor quality particles, all subsequent processing was carried out in in RELION-3.0^45^. Particles were re-extracted using a 432 x 432 pixel box from micrographs that had been re-processed using the MotionCor2^46^ implementation in RELION-3.0, with CTF estimation by CTFFIND4^47^.

Initial processing of the Triton-X100 extracted particles produced 2D classes with close to top down views that allowed preliminary counting of the subunits around the perimeter of the object, although the lack of purely top down views prevented unambiguous assignment. 27435 particles were selected after classification and used to generate *ab initio* initial models with C33 and C34 symmetry. 3D classification was carried out with C33 and C34 symmetries applied and the C33 job produced a class containing 15634 particles that refined to 3.8 Å using gold standard refinement. Reclassification of the original particles produced a 19520 particle set that led to a 3.1 Å map following Bayesian polishing^48^ and per-particle CTF refinement. This allowed *de novo* model building of the RBM3/β-collar domains (residues 231-438) but all other regions of the map remained untraceable. Attempts at reconstructing with lower symmetry were hindered by the low particle number.

A larger, DDM-extracted, dataset was collected that contained 188007 particles after 2D classification. 3D classification applying C33 symmetry resulted in one good class with 106745 particles which were then used in a C1 symmetry refinement. This produced density in the ring below the C33 ring with a clear periodicity that could be counted as C21. Refinement of this particle set with the common symmetry of C3 applied produced a 3.3 Å map, following Bayesian polishing and CTF refinement, that revealed an RBM fold in the 21-fold symmetric ring. However, the quality of this portion of the map was not sufficiently detailed to allow *de novo* model building. As the proportion of particles that produced a sub-3.5 Å map were similar between the two different detergent extractions, and the maps produced were indistinguishable, we created a combined dataset containing the post-2D classification particles from the Triton-X100 and DDM extractions and a small dataset from a DDM extracted sample that had been exchanged into amphipol A8-35. This dataset, containing 273493 particles was subjected to 3D classification applying C3 symmetry, using the DDM-only model low pass filtered as a reference. After two rounds of 3D classification, two good classes were produced, containing 126285 and 59163 particles. The first of these classes refined to a pure C33 object in the RBM3 region, but the second class produced a map with ∼11.3 subunits per “asymmetric unit” in the C3 symmetry. We therefore re-refined this class applying C34 symmetry, which produced a 3.3 Å gold standard map. Refinement of the “C34” particles in C1 produced periodicity in the ring below the RBM3 ring consistent with C22 symmetry.

Due to the increased complexity of the sample, we collected a large A8-35 exchanged dataset and created a composite Triton-X100/DDM/A8-35 dataset containing 449142 particles after 2D classification. These particles were then subjected to a supervised 3D classification in C1, using C33 and C34 maps as references, producing classes with 308536 and 140606 particles respectively. The C33 class was subjected to a further round of classification, producing a good class with 175233 particles that was refined in C3 to an overall resolution of 2.9 Å following Bayesian polishing and CTF refinement. Further focused classification and refinement of the C33 particles with a mask around the RBM3/β-collar region, and with C33 symmetry applied, produced a 2.6 Å map from 77849 particles. Further focused classification and refinement of the C33 particles with a mask around the RBM2_inner_ region, and with C21 symmetry applied, produced a 2.9 Å map from 84797 particles. Attempts to improve the resolution of the RBM2_outer_/RBM1 region through particle subtraction, multi-body refinement and local averaging were unsuccessful. Initial refinements of the entire object produced maps with nine strong copies of the RBM2_outer_/RBM1 pair and weaker density in the gaps between copies 3 and 4, 6 and 7 and 9 and 1. This weaker density was consistent with being a superposition of two copies of the RBM2_outer_/RBM1 density. However, the spacing of these domains was such that these gaps could not accommodate a full RBM2_outer_/RBM1 pair without structural rearrangement, and we reasoned that the density observed could be produced by rotational misalignment of a subset of the particles producing “ghost” density from the strong domains. In order to test this, we masked around the nine strong domain pairs and used this mask in a focused refinement, with the logic that if extra copies were genuinely ordered they would appear in the final, unmasked, map. This was not found to be the case. The C34 class was refined with C2 symmetry applied and produced a 3.3 Å map after Bayesian polishing and CTF refinement. Further focused refinement of the C34 particles with a mask around the RBM3/β-collar region, and with C34 symmetry applied, produced a 2.8 Å map. Further focused classification and refinement of the C34 particles with a mask around the RBM2_inner_ region, and with C22 symmetry applied, produced a 3.1 Å map from 87107 particles. Similar analysis of the RBM2_outer_/RBM1 region was applied as in the C33 refinements, with similar results, but in this case ten copies of the domain pair could be placed. All processing statistics are summarised in Extended Data Tables 1 and 2.

### Model building and refinement

A monomer model for the RBM3 and the β-collar (residues 231-438) was built manually in Coot^49^ using the 2.6 Å map with C33 symmetry applied, assembled into a 33-fold model, and refined using phenix.real_space_refine^50^. A monomer model for RBM2 (residues 125-222) was built manually in Coot using the 2.9 Å map with C21 symmetry applied, assembled into a 21mer of the RBM2_inner_ region and refined using phenix.real_space_refine. The whole 33mer was assembled from these two structures in the 2.9 Å map with C3 symmetry applied. The two main rings were joined by manually building the linkers in Coot. Nine copies of the high resolution RBM2 domain were placed manually in the RBM_outer_ domain densities of a 4 Å lowpass filtered version of the C3 map, and rigid body refined. Nine copies of a RaptorX generated homology model of RBM1 (residues 50-106) were manually positioned in the density underneath the RBM_outer_ domains and rigid body refined. The completed 33mer was refined with phenix.real_space_refine, using the higher resolution C33 and C21 structures as reference models. The RBM3/β-collar monomer built in the C33 map was used to assemble a 34-fold model in the 2.8 Å map with C34 symmetry applied, and was refined using phenix.real_space_refine. A 22-fold RBM2_inner_ model was assembled in the 3.1 Å map with C22 symmetry applied, using the RBM2 monomer built in the C21 map, and was refined using phenix.real_space_refine. The whole 34mer was assembled from these two structures in the 3.3 Å map with C2 symmetry applied. The two main rings were joined by manually building the linkers in Coot. Ten copies of the RBM_outer_/RBM1 domain pairs from the 33mer model were placed manually in the appropriate densities of a 4 Å lowpass filtered version of the C2 map, and were rigid body refined. The completed 34mer was refined with phenix.real_space_refine, using the higher resolution C34 and C22 structures as reference models. All models were validated using Molprobity^51^. All refinement and validation statistics are summarised in Extended Data Tables 1 and 2. Conservation analysis was carried out using the Consurf server^52^. Figures were prepared using Pymol (The PyMOL Molecular Graphics System, Version 2.0 Schrödinger, LLC) and ChimeraX^53^.

## Author Contributions

SJ & SML designed the project, interpreted the data and wrote the first draft of the paper. SJ analysed the data. YHF cloned, expressed and purified protein samples, and made and optimised EM grids. JCD made and screened grids. JCD & SML collected the EM data. EF expressed and purified samples and made EM grids. LK made constructs and performed preliminary purification experiments. All authors commented on drafts of the manuscript.

