## Extended Data for "Structure of the bacterial flagellar rotor MS-ring: a minimum inventory/maximum diversity system"

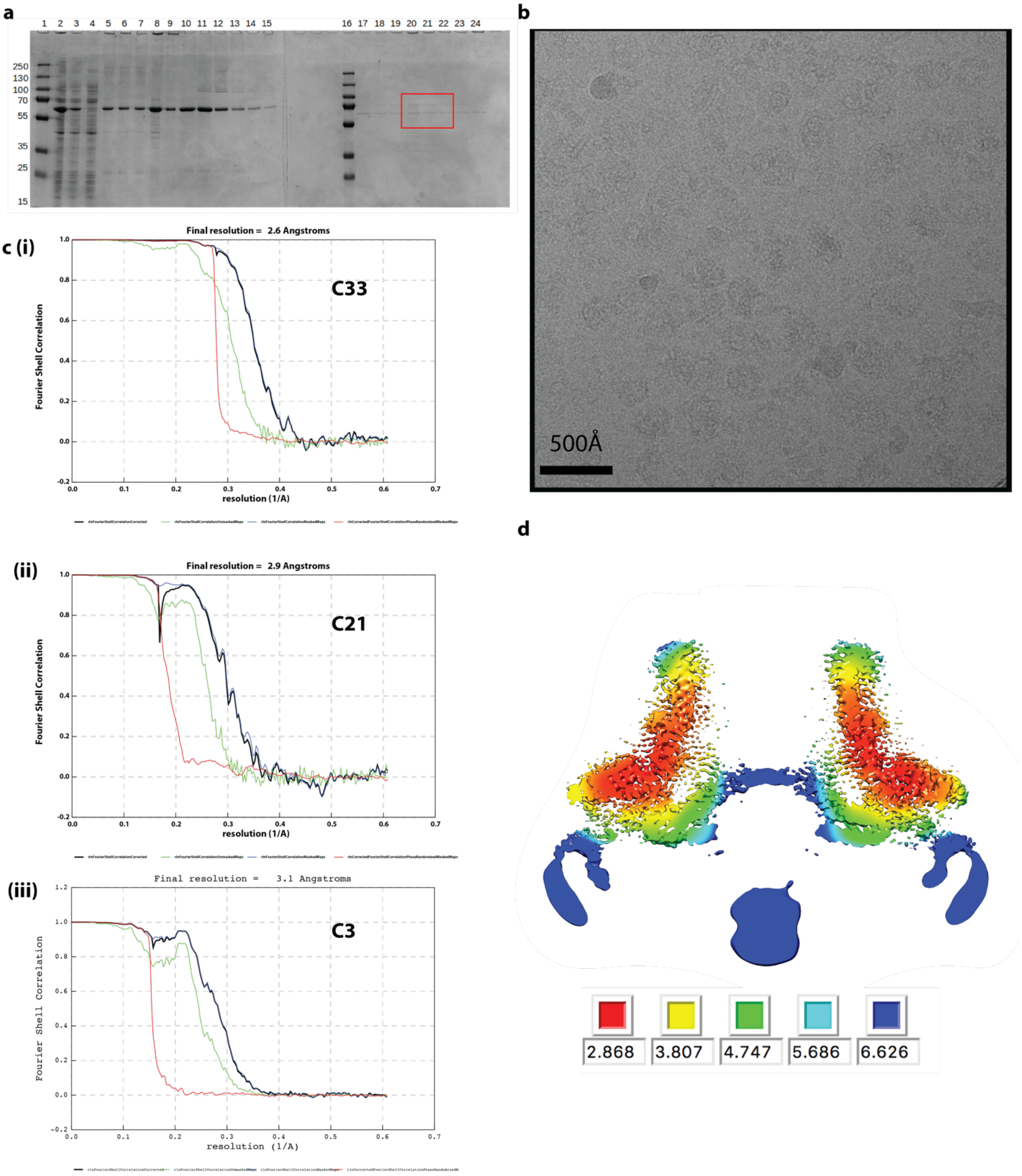

**Extended Data Figure 1**. **a**. SDS-PAGE of samples taken throughout purification of FliF in DDM. Lanes contain (1, 16) PageRuler Markers (2) whole cell lysate (3) supernatant from low speed spin (4) supernatant from high speed spin (5) solubilised membranes (6) supernatant from second low speed spin (7) supernatant from second high speed spin (8) resuspended pellet from second high speed spin (9-15 and 17-20) fractions from top to bottom of sucrose gradient post-high speed equilibration. Note – this gel shows samples from sucrose gradient without glutaraldehyde run in parallel with tubes containing glutaraldehyde. The fractions equivalent to those indicated (red box) were selected from the cross-linked gradients and used for structural analysis. **b**. Example micrograph (1.5 µm defocus) of cross-linked FliF on a graphene oxide surface. **c.** FSC curves from PostProcessing in RELIONv3.0 for volumes calculated in (i) C33, (ii) C21 and (iii) C3 respectively. **d**. Slab through C3 volume coloured by local resolution as estimated using RELIONv3.0.

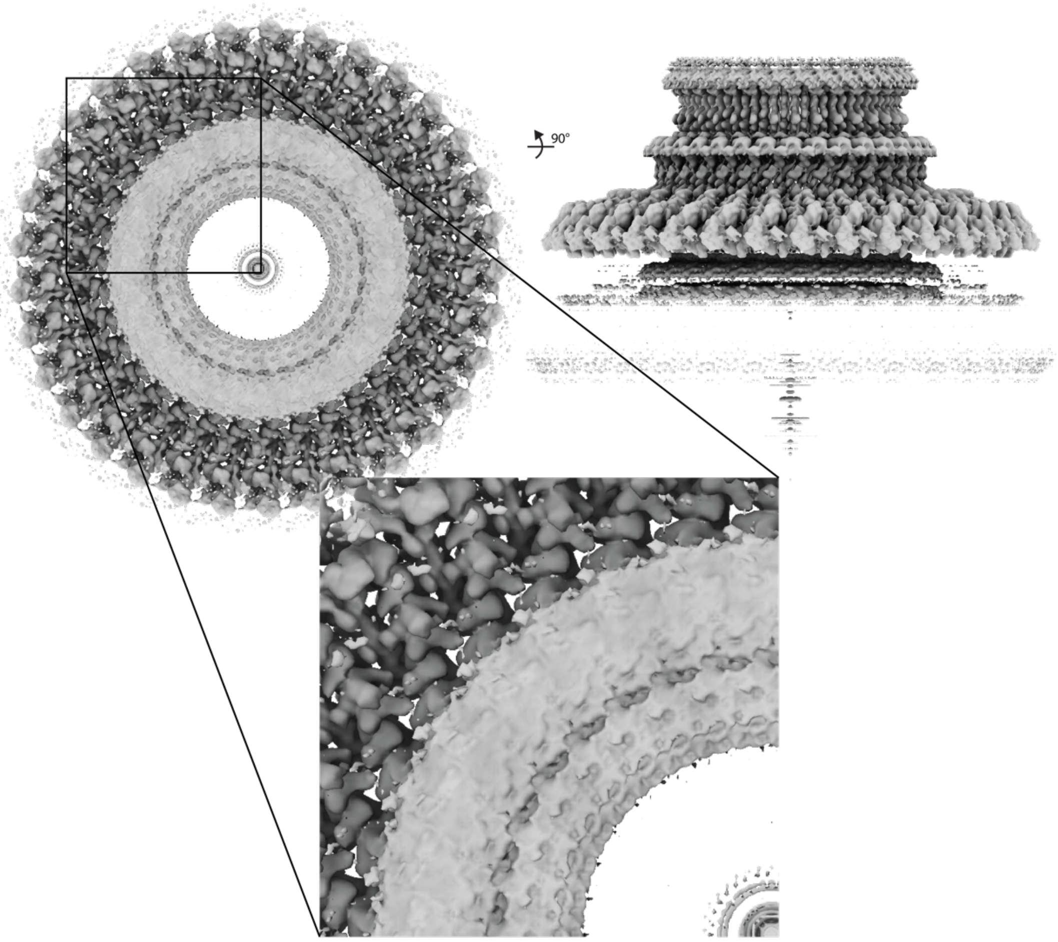

**Extended Data Figure 2.** Volume generated by refinement in C33 shows lack of detail in $RBM2_{inner}$ region below, later explained by C21 symmetry in this region.

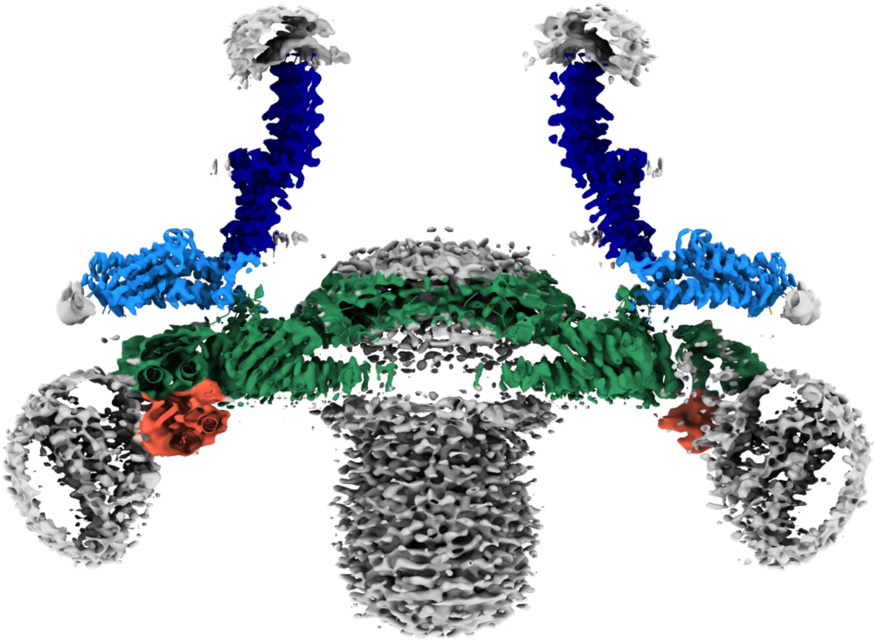

**Extended Data Figure 3.** Slab through central section of composite C33/C21/C3 volume reveals layered density derived from the detergent micelle at the periphery of the RBM3 ring and a central column of density below the C21 ring that presumably results from density associated with the 24 copies of RBM1 that are not located elsewhere in the map, the N-terminal trans-membrane helices attached to these and associated detergent.

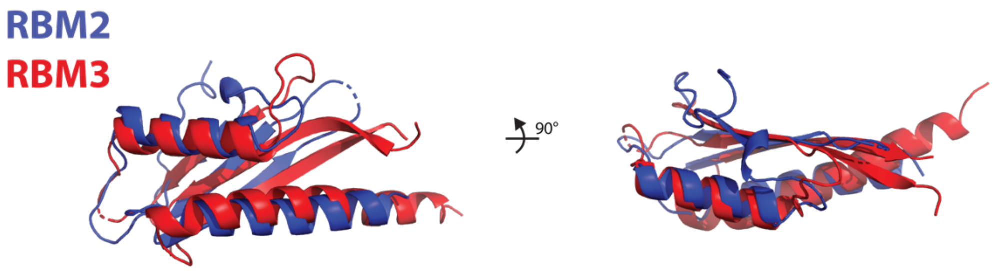

**Extended Data Figure 4.** Overlay of RBM2 and RBM3 domains of FliF (chain A)

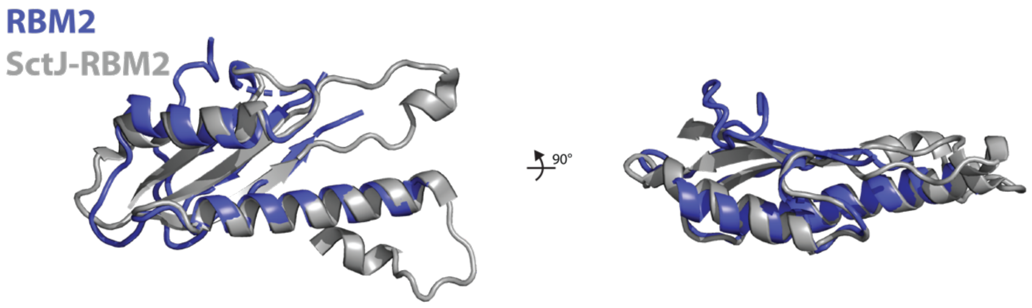

**Extended Data Figure 5.** Superposition of the RBM2 domain of FliF on the closest structural homologue – the RBM2 of SctJ

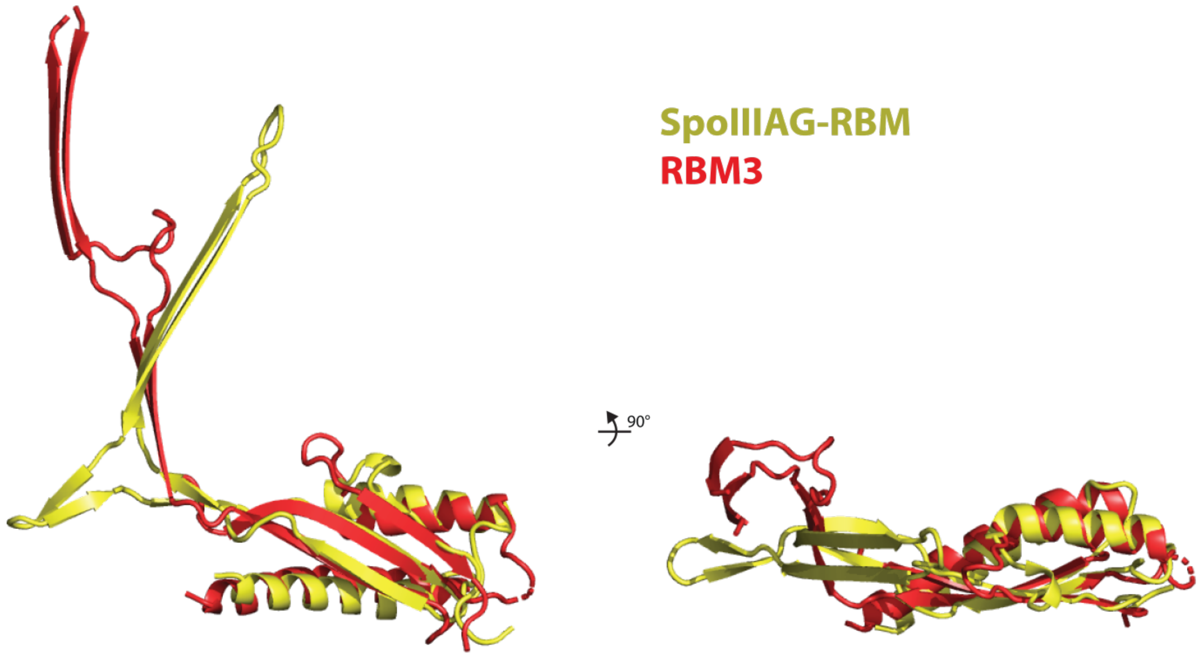

**Extended Data Figure 6.** Superposition of the RBM3 domain of FliF on the SpoIIIAG RBM domain. The beta-insertions are not used to derive the superposition and the different relationship between the RBM domains and these inserts can therefore be appreciated.

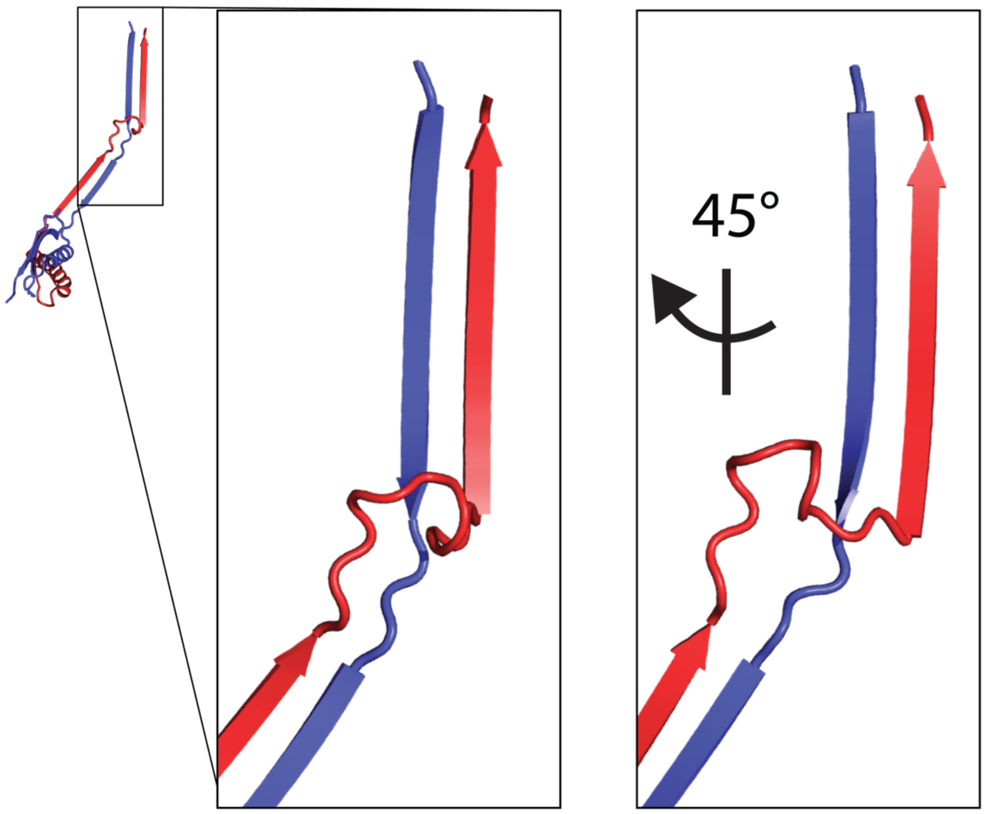

**Extended Data Figure 7.** The N-terminal (blue) and C-terminal strands (red) of the beta-insert cross at the transition between tilted and vertically oriented strands.

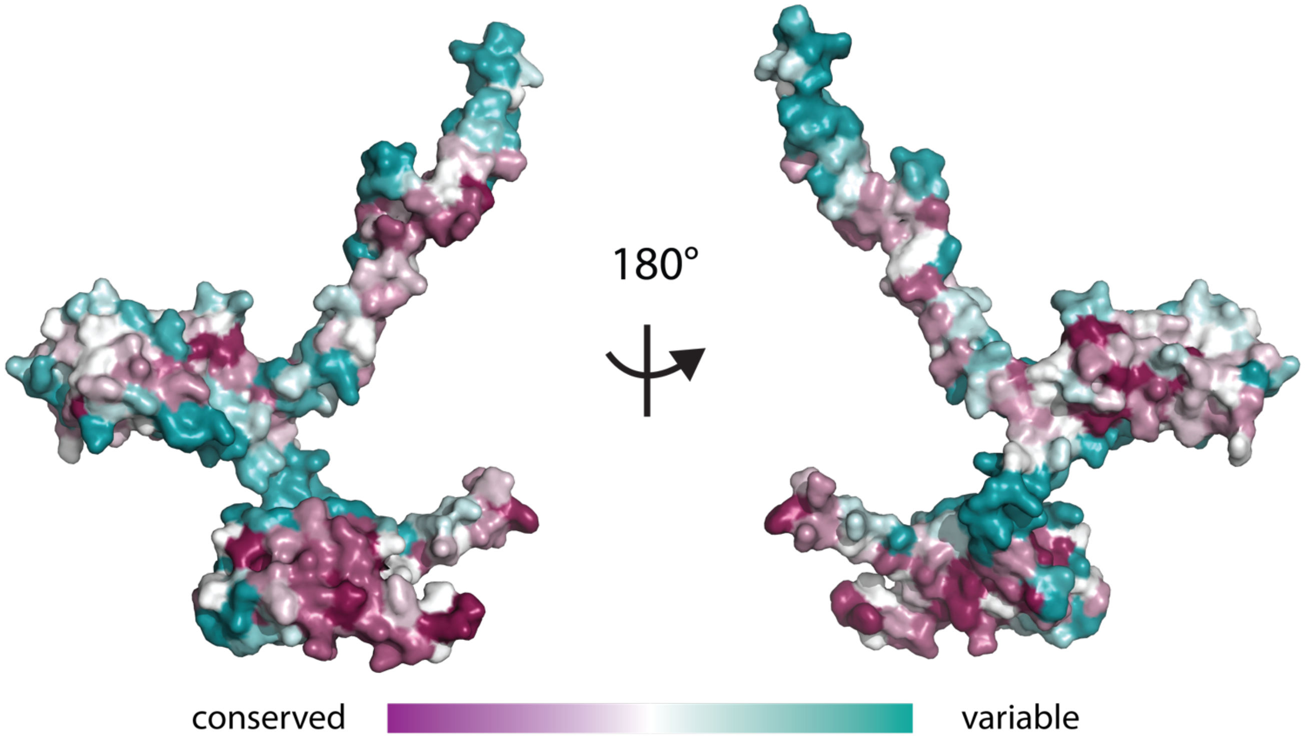

**Extended Data Figure 8.** Conservation of FliF between bacterial species is plotted on the surface of the monomer (ConSurf http://consurf.tau.ac.il) revealing that the monomer-monomer interfaces are the most conserved regions.

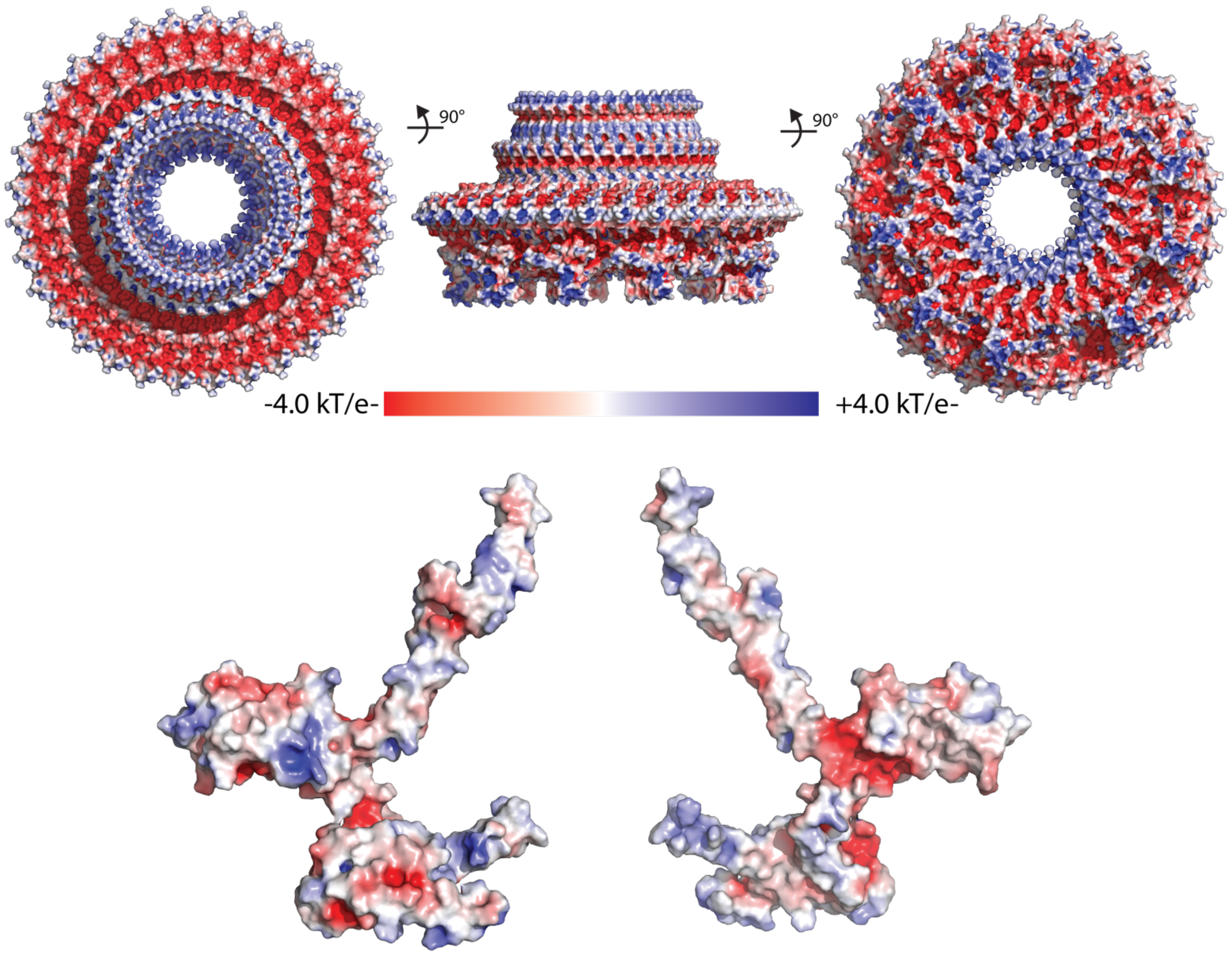

**Extended Data Figure 9.** The Electrostatic potential is mapped onto the surface of the FliF assembly (upper panel) and monomer using APBS within PyMol, revealing that the overall object is highly charged whilst the monomer interfaces are largely hydrophobic.

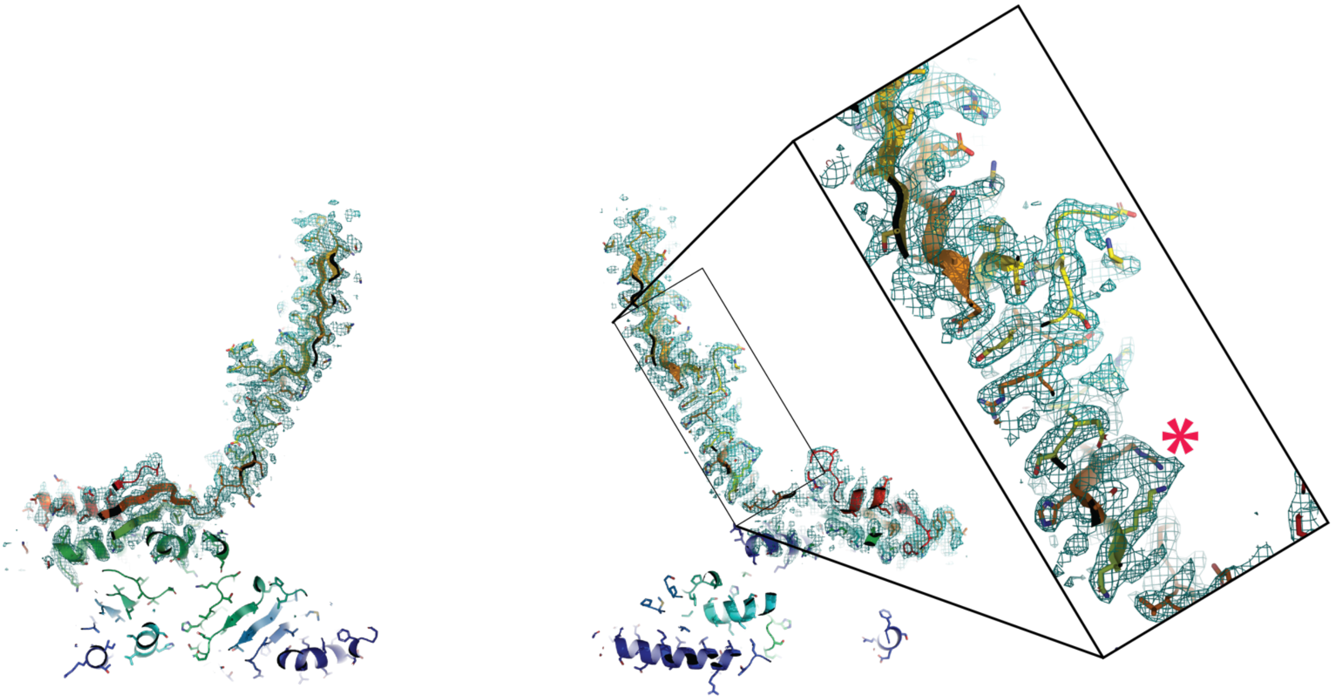

**Extended Data Figure 10.** A putative glutaraldehyde cross-link (marked with an asterisk) is observed in the beta-collar region of FliF.

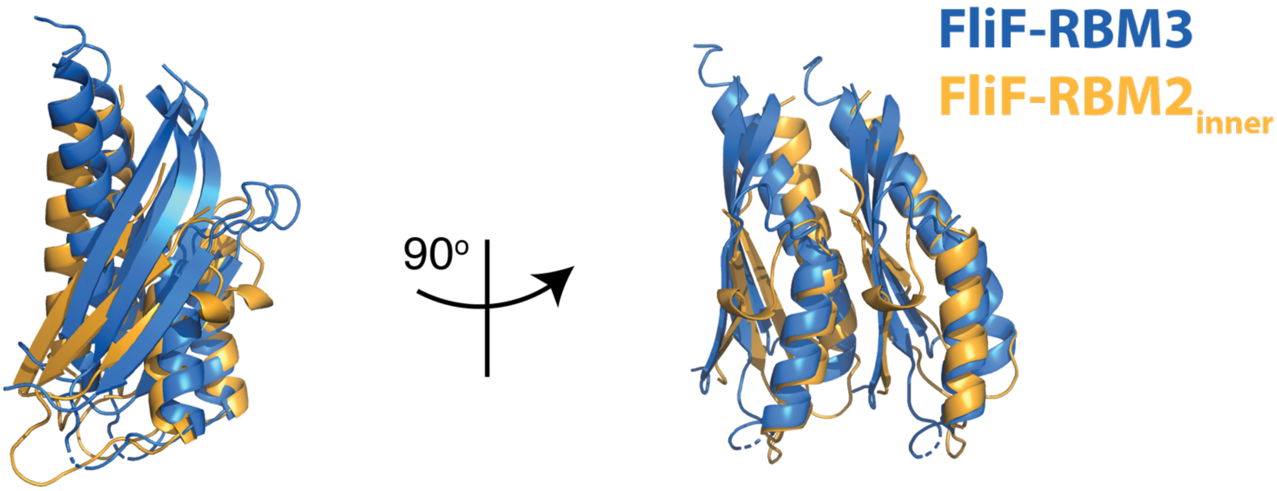

**Extended Data Figure 11.** Superposing a pair of neighbouring RBM3 domains on to a pair of neighbouring RBM2inner by aligning the first domain shows the rearrangements driven by the C33 versus C21 packing.

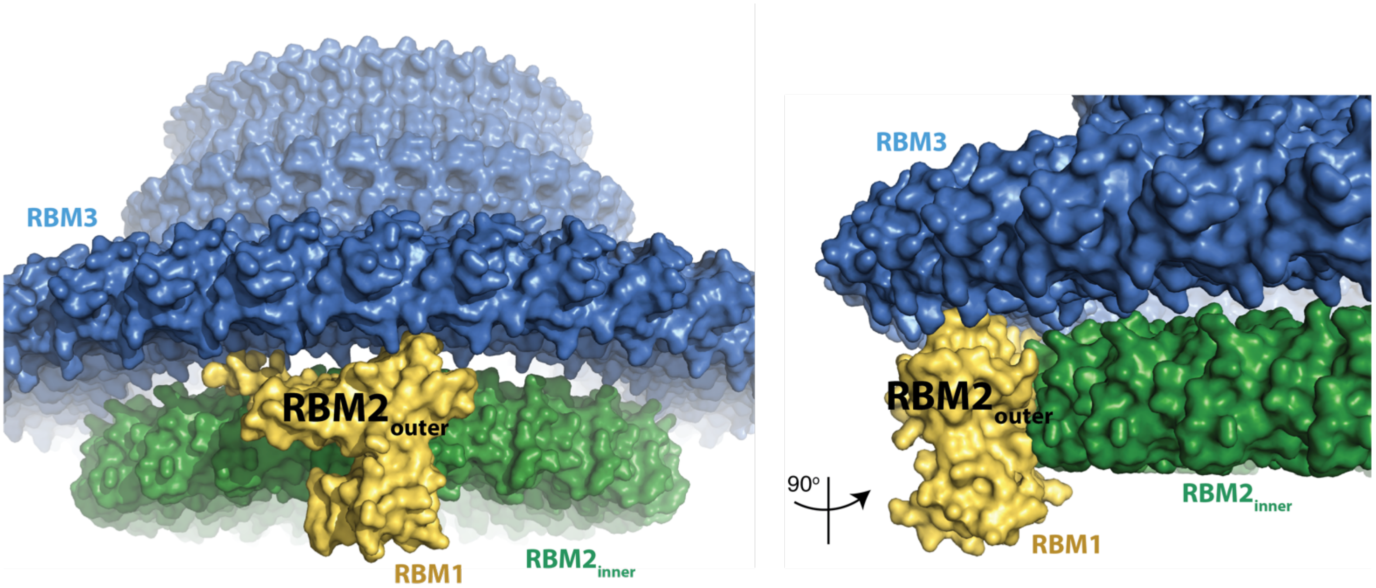

**Extended Data Figure 12.** The RBM2$_{outer}$ domains provide the major contact between the RBM2$_{inner}$ and RBM3 rings adapting between the C21 and C33 symmetries.

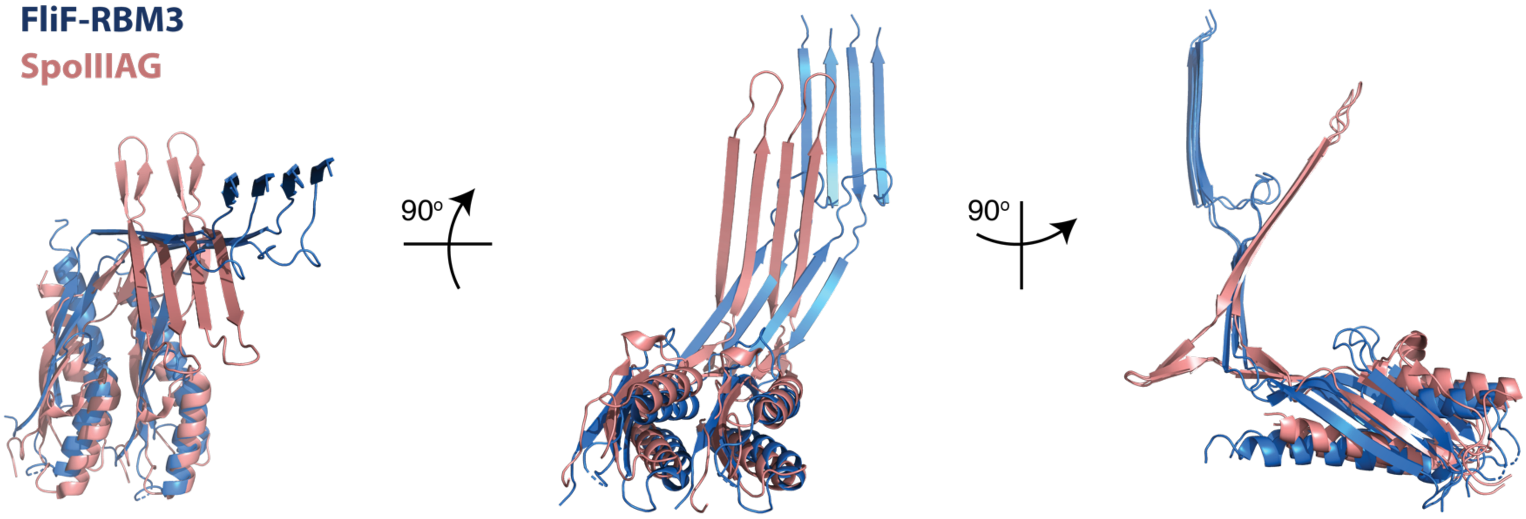

**Extended Data Figure 13.** Superposing a pair of neighbouring RBM3 domains on to a pair of neighbouring SpoIIAG RBM domains by aligning the first domain shows the subtle alteration in packing needed to form the C33 rather than C30 assemblies.

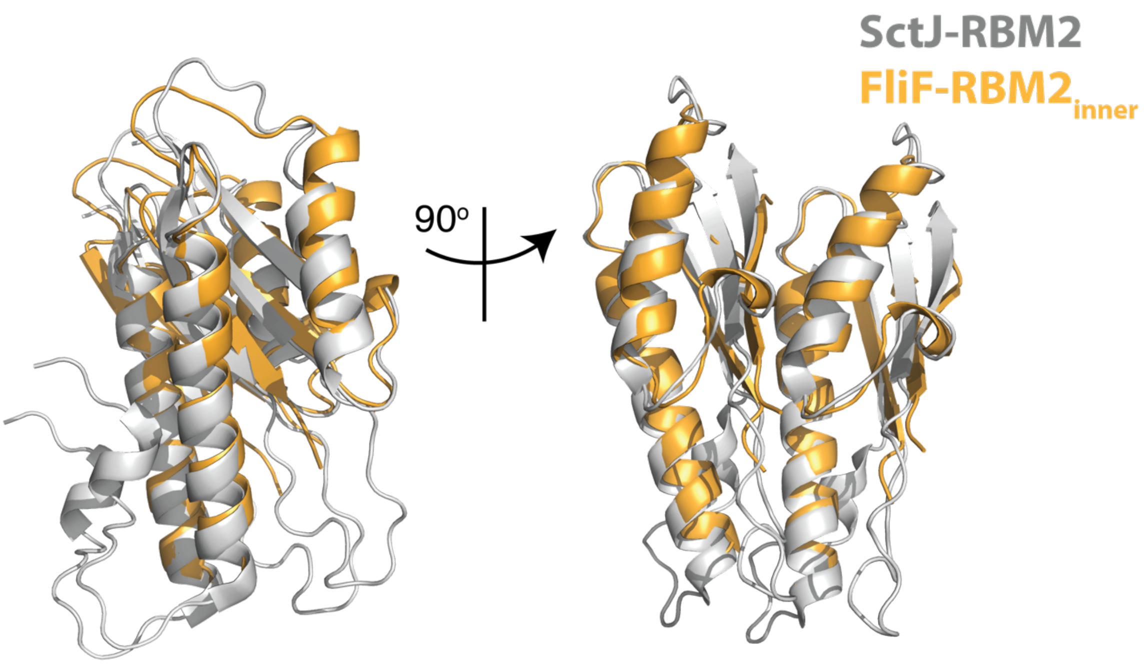

**Extended Data Figure 14.** Superposing a pair of neighbouring $RBM2_{inner}$ domains on to a pair of neighbouring SctJ RBM2 domains by aligning the first domain shows the subtle alteration in packing needed to form the C21 rather than C24 assemblies.

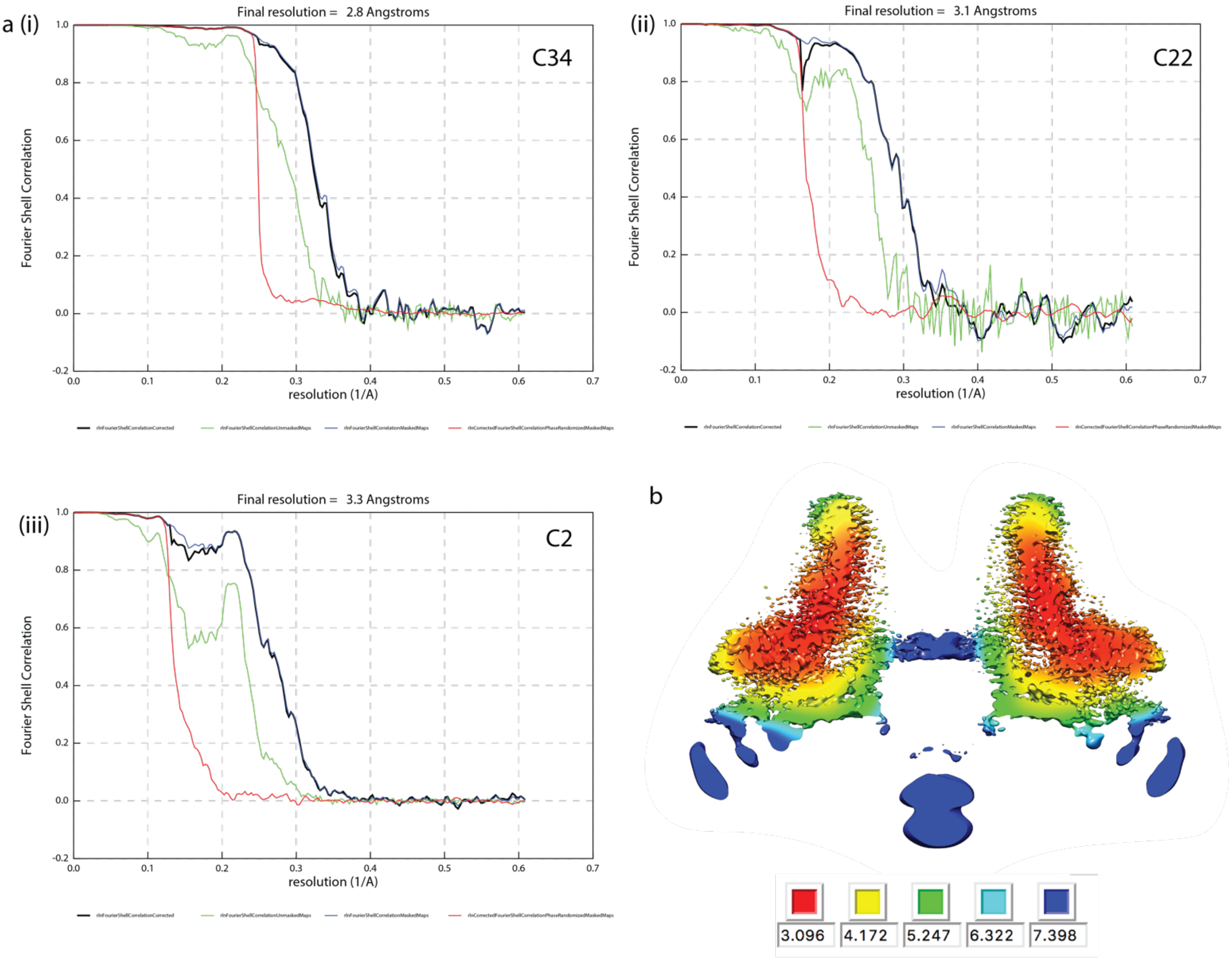

**Extended Data Figure 15.** **a.**FSC curves from PostProcessing in RELIONv3.0 for volumes calculated in (i) C34, (ii) C22 and (iii) C2 respectively. **b.** Slab through C2 volume coloured by local resolution as estimated using RELIONv3.0.

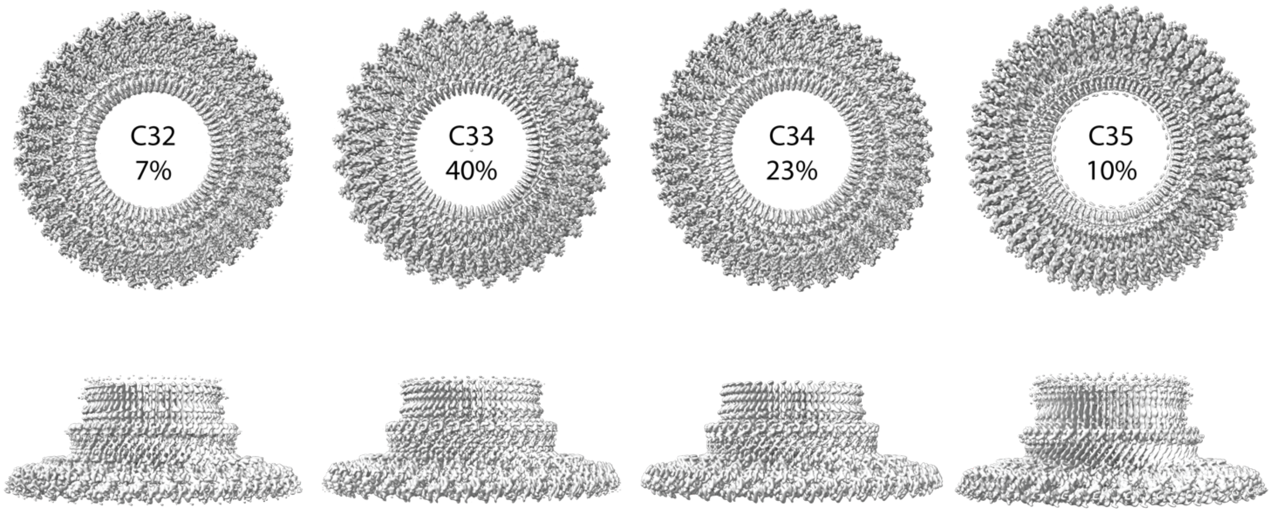

**Extended Data Figure 16.** Distribution of particles between different symmetries in the RBM3 ring/β-collar region following supervised 3D-classification. 20% of particles were allocated to a C36 class, but the volume was uninterpretable from this class and presumably reflected damaged particles / particles with a variety of other symmetries.

**Extended Data Table 1: 33-fold FliF Cryo-EM data collection, refinement and validation statistics**

| | 33-fold FliF<br>Whole map<br>(EMD-10143)<br>(PDB-6SCN) | 33-fold FliF<br>RBM3/β-collar<br>(EMD-10145)<br>(PDB-6SD1) | 33-fold FliF<br>$RBM2_{inner}$<br>(EMD-10146)<br>(PDB-6SD2) |
|---|---|---|---|
| **Data collection and processing** | | | |
| Magnification | 165,000 | 165,000 | 165,000 |
| Voltage (kV) | 300 | 300 | 300 |
| Electron exposure (e–/Å$^2$) | 48 | 48 | 48 |
| Defocus range (μm) | 0.5–4 | 0.5–4 | 0.5–4 |
| Pixel size (Å) | 0.822 | 0.822 | 0.822 |
| Symmetry imposed | C3 | C33 | C21 |
| Initial particle images (no.)* | 449142 | 449142 | 449142 |
| Final particle images (no.) | 175233 | 77849 | 84797 |
| Map resolution (Å) | 3.1 | 2.6 | 2.9 |
| FSC threshold | 0.143 | 0.143 | 0.143 |
| Map resolution range (Å) | 2.9-6.6 | 2.5-3.1 | 2.9-3.6 |
| **Refinement** | | | |
| Initial model used (PDB code) | *Ab initio* | *Ab initio* | *Ab initio* |
| Model resolution (Å) | 3.1 | 2.6 | 2.9 |
| FSC threshold | 0.143 | 0.143 | 0.143 |
| Model resolution range (Å) | | | |
| Map sharpening *B* factor (Å$^2$) | -72 | -62 | -104 |
| Model composition | | | |
| Non-hydrogen atoms | 60582 | 39377 | 13566 |
| Protein residues | 7875 | 4984 | 1848 |
| Ligands | 0 | 0 | 0 |
| *B* factors (Å$^2$) | | | |
| Protein | 130 | 53 | 59 |
| Ligand | N/A | N/A | N/A |
| R.m.s. deviations | | | |
| Bond lengths (Å) | 0.007 | 0.003 | 0.009 |
| Bond angles (°) | 0.817 | 0.442 | 0.770 |
| Validation | | | |
| MolProbity score | 2.0 | 1.7 | 2.4 |
| Clashscore | 13.0 | 7.1 | 6.7 |
| Poor rotamers (%) | 0.8 | 0.0 | 5.6 |
| Ramachandran plot | | | |
| Favored (%) | 94.3 | 95.7 | 92.7 |
| Allowed (%) | 4.9 | 4.3 | 7.3 |
| Disallowed (%) | 0.8 | 0.0 | 0.0 |

* Particle numbers quoted are post-2D clean-up

**Extended Data Table 2: 34-fold FliF Cryo-EM data collection, refinement and validation statistics**

| | 34-fold FliF Whole map (EMD-10147) (PDB-6SD3) | 34-fold FliF RBM3/β-collar (EMD-10148) (PDB-6SD4) | 34-fold FliF $RBM2_{inner}$ (EMD-10149) (PDB-6SD5) |
|---|---|---|---|
| **Data collection and processing** | | | |
| Magnification | 165,000 | 165,000 | 165,000 |
| Voltage (kV) | 300 | 300 | 300 |
| Electron exposure (e–/Å$^2$) | 48 | 48 | 48 |
| Defocus range (μm) | 0.5–4 | 0.5–4 | 0.5–4 |
| Pixel size (Å) | 0.822 | 0.822 | 0.822 |
| Symmetry imposed | C2 | C34 | C22 |
| Initial particle images (no.)* | 449142 | 449142 | 449142 |
| Final particle images (no.) | 140606 | 140606 | 87107 |
| Map resolution (Å) | 3.3 | 2.8 | 3.1 |
| FSC threshold | 0.143 | 0.143 | 0.143 |
| Map resolution range (Å) | 3.1-7.4 | 2.7-3.6 | 3.0-3.9 |
| **Refinement** | | | |
| Initial model used (PDB code) | EMD-10143 | EMD-10143 | EMD-10143 |
| Model resolution (Å) | 3.3 | 2.8 | 3.1 |
| FSC threshold | 0.143 | 0.143 | 0.143 |
| Model resolution range (Å) | | | |
| Map sharpening *B* factor (Å$^2$) | -67 | -85 | -103 |
| Model composition | | | |
| Non-hydrogen atoms | 63144 | 40562 | 14212 |
| Protein residues | 8212 | 5134 | 1936 |
| Ligands | 0 | 0 | 0 |
| *B* factors (Å$^2$) | | | |
| Protein | 139 | 41 | 51 |
| Ligand | N/A | N/A | N/A |
| R.m.s. deviations | | | |
| Bond lengths (Å) | 0.008 | 0.009 | 0.006 |
| Bond angles (°) | 0.824 | 0.657 | 0.617 |
| Validation | | | |
| MolProbity score | 2.3 | 1.9 | 2.0 |
| Clashscore | 12.3 | 5.7 | 7.1 |
| Poor rotamers (%) | 1.2 | 0.8 | 1.4 |
| Ramachandran plot | | | |
| Favored (%) | 88.5 | 89.7 | 91.7 |
| Allowed (%) | 10.8 | 9.6 | 8.3 |
| Disallowed (%) | 0.7 | 0.7 | 0.0 |

* Particle numbers quoted are post-2D
